# Degradation of Visible Autumn Icons and Conservation Opportunities: Trends in Deciduous Forest Loss in the Contiguous US

**DOI:** 10.1101/2021.03.29.437570

**Authors:** L. M. Dreiss, J.W. Malcom

## Abstract

Temperate deciduous forests are one of the most visible biomes on Earth because of their autumn aesthetics and because they harbor some of the most heavily populated regions. Their ability to attract visitors may increase opportunities for people to experience nature, which has been linked to greater conservation action. Identifying regions with high leaf-peeping opportunities and regions where color has been lost to landscape conversion may help to inform these connections. We use spatial overlay analyses to quantify temperate deciduous forest coverage, disturbance, and protections in each U.S. ecoregion. We evaluated recent (1984-2016) and predicted (2016-2050) disturbance under extreme future scenarios. Almost all ecoregions saw a decline in deciduous forest cover between 1985 and 2016. Some ecoregions with the greatest opportunities for leaf-peeping are also underrepresented in the protected areas network and vulnerable to additional losses. Under economic-growth forecasting scenarios, losses are predicted to continue. However, environmentally focused scenarios suggest there is still opportunity to reverse deciduous forest loss in some ecoregions. Differences in forest loss between predictions scenarios emphasize the importance of human approaches in securing environmental stability. Increasing public exposure to temperate forests may help ensure conservation of more natural areas and preserve the quantity and quality of autumn forest viewing.

**Key Points:**

- Temperate deciduous forests aesthetics attract visitors to experience nature, but degradation and loss can hinder connections.
- US ecoregions with the greatest leaf-peeping opportunities are underrepresented in the protected areas and vulnerable to additional losses.
- Differences in predictions scenarios emphasize the importance of conservation action, which may be linked to human connections with nature.

## 1 Introduction

Accelerating landscape change threatens biodiversity and climate stability worldwide, with significant implications for society through a degradation of nature’s benefits to people (i.e., “ecosystem services”; Carvahlo and Szlafsztein, 2019; Colvin et al., 2019; Diaz et al., 2020; Gómez-Baggethun et al., 2019; IPBES, 2019; Sweeney et al., 2004). The science makes clear that transformative action is needed to address the threats to nature, with major initiatives being developed at national and international levels (e.g., Convention on Biological Diversity, 2020; Udall, 2020). While preservation and restoration are common strategies for addressing these crises (e.g., Clancy et al., 2020; Dinerstein et al., 2019, 2020; Minin et al., 2017; Peng et al., 2019; Scolozzi et al., 2014), approaches may also include increasing opportunities for people to experience natural environments: when people connect with nature they are also more likely to act in ways that benefit the Earth (Ives et al., 2016). Axiological relationships are often tied to feelings of duty and stewardship and previous research suggests a similar connection between a person’s bond with nature and the probability of them taking conservation action (Cooper et al., 2016). Connections with nature can happen emotionally, physically, intellectually, and spiritually, but most often, through direct experience (Wang et al., 2016). Identifying highly visible environments and understanding past and future changes that people have or will experience is therefore important to understanding one social dimension of conservation.

Although they only cover ~7.5% of Earth’s terrestrial land surface, temperate forests harbor exceptional biodiversity and carbon stores (Hofmeister et al., 2019; Keith et al., 2009; Pan et al., 2011; Thurner et al., 2013) and provide key ecosystem services such as water filtration (Brandt et al., 2014; DellaSala et al., 2011). Further, temperate forests are one of the most visible biomes because heavily populated and developed regions, including in the United States, are located in or near the biome (Haddad et al., 2015). At the same time, a long history of settlement and development has had dramatic impacts on the forest ecosystems and their biological diversity (e.g., Gunn et al., 2019; Pennington et al., 2010; Thompson and Jones, 1999). Although most of the major land changes may have occurred in the past, more recent anthropogenic landscape modifications mean continued forest loss, biodiversity loss, and degradation of ecosystem services. Understanding recent and predicted temperate forest conversion can help identify spatial patterns and clarify the critical role of human perceptions of change that may affect forest conservation.

Among the ecosystem services provided by temperate forests, the biome provides a unique aesthetic appeal with economic benefits. About 15% of the tree species of the temperate regions of the world change their leaf color from green to yellow or red in autumn, a percentage that can reach 70% of species in some regions of the US (Archetti et al., 2013). In this respect, deciduous forests are very visible cultural icons. Autumn aesthetic is often described in terms of its “complex”, “reassuring”, and “soothing” qualities, attracting outsiders seeking beauty and relaxation in special landscapes (Eroglu and Demir, 2016). As such, many U.S. cultures and economies rely on autumn traditions and ‘leaf-peeping’ tourism, which contributes billions of dollars each year - up to a quarter of annual tourism profit - to state economies of the eastern US. (Sandifer et al., 2015). Continued landscape conversion threatens to reduce the autumn color display directly (i.e., tree removal) or indirectly through increased forest stress, and introduces concerns for reduced future aesthetic values and tourism revenues. Ultimately, fewer visitors to these forests also means fewer opportunities to connect people and nature.

Given the current global biodiversity and climate crises, more conservation action is needed and there is much to gain with temperate forests as a connection between people and nature. The reverse - using personal connections with nature to specifically emphasize conservation of temperate forests - could also result in major contributions to biodiversity conservation and climate mitigation. In the U.S., temperate forests are home to high species endemism and hotspots of imperiled species biodiversity (Rosa and Malcom, 2020). Additionally, intact temperate forests remove sufficient atmospheric CO_2_ to reduce national annual net emissions by 11% and may be a significant contributor to global climate stabilization (Dinerstein et al., 2020; Moomaw et al., 2019). Studies suggest that U.S. temperate forests are likely to act as carbon sinks for decades to come (Finzi et al., 2020). However, most remaining examples of natural ecosystems are fragmented and highly modified with intensive human activities, posing continued threats to biota, carbon sequestration, and ecosystem aesthetics.

General trends in temperate forest land use emphasize continuing rural residential development largely driven by aesthetics, climate and access to recreation. Beyond the direct impacts of forest loss and expanding anthropogenic land cover, biodiversity and ecosystem functions are likely to suffer from fragmentation; forests are smaller, more isolated, and have a greater area located near the edge. This includes degraded ecosystem processes, reduced species diversity, and changes in species all of which affect autumn foliage viewing quality (Haddad et al., 2015; Zhang et al., 2017; Zhang et al., 2019). Fragmentation of the forested landscape may also contribute to the degradation of autumn foliage viewing in a way that is less apparent than with outright forest loss: viewers may fail to notice the creeping change in viewing quality until it is too late (i.e., boiling frog theory; Soga and Gaston, 2018).

Given the importance of temperate forests as a symbol for nature and its beauty, forest conversion is among the most distinct impacts of human activity on the forest environment and aesthetic. Here, we identify to what extent each U.S. ecoregion is characterized by deciduous forest and quantify recent (1984-2016) and predicted (2016-2050) forest disturbance. The ability to identify regions with high leaf-peeping opportunities and regions where color has been lost to landscape conversion will help to inform connections between people and the forested landscape. Additionally, understanding patterns of temperate forest loss is also critical to guide efforts in biodiversity conservation and climate mitigation: new calls to expand the protected areas network will benefit from this and subsequent knowledge that builds upon these findings.

## 2 Materials and Methods

We used spatial overlay analyses to estimate temperate forest losses and predicted changes across the U.S., summarized by EPA Level 2 ecoregions (EPA, 2006). Two main datasets were used for these calculations. First, the National Land Cover Database (NLCD, MRLC) uses digital change detection methods to identify land cover, changes and trends for the United States (Homer et al., 2020). The recently released NLCD 2016 database offers improved cyclical updating of U.S. land cover and associated changes, greatly advancing large-area land cover monitoring through an updated suite of products (Yang et al., 2018). Forest disturbance date is a key addition that provides opportunities to examine US forest cover change patterns from 1985-2016. This product combines information from the NLCD 2016 change detection, land cover classification, and the LANDFIRE Vegetation Change Tracker (VCT) disturbance product to assess where disturbance occurred for forest areas every 2-3 year interval.

NLCD classifications for deciduous forest and mixed forest were included in areal calculations for available years between 1992 - 2016 (1992, 2001, 2004, 2006, 2008, 2011, 2013, and 2016). Modelled historical land use and land cover for the contiguous US were used for estimating forest cover prior to 1992 (Sohl et al., 2018). Similarly, all forest change identified in the forest disturbance date dataset between 1985-2016 that occurred on deciduous and mixed forest land cover types was used in calculations of forest loss for each ecoregion.

Second, we used spatially explicit predictive models of 2050 land cover to estimate future forest losses (Sohl et al., 2014). The most extreme forecasting scenarios which were developed based on characteristics consistent with the Intergovernmental Panel on Climate Change (IPCC) Special Report on Emission Scenarios were used to understand the variability in predictions. The economic-growth scenario assumes rapid economic development and very high population growth globally (15 billion by 2100). Initial estimates of this scenario suggest over 500,000 km^2^ of deciduous/mixed forests would be lost in the contiguous US between 2005 and 2100 (Sohl et al., 2014). The sustainability scenario reflects an emphasis on environmental protection and social equity with lower rates of global human population growth and intermediate levels of economic development. In contrast, over 59,000 km2 of forests may be gained between 2005 and 2100 under the environmental protection scenario (Sohl et al., 2014).

Data on protected areas are from the PADUS 2.0 database. We use U.S. Geological Survey’s Gap Analysis Program (GAP) codes, which are specific to the management intent to conserve biodiversity. GAP 1 and 2 areas are managed in ways typically consistent with conservation and are considered ‘protected’ in this context. We used ArcGIS v. 10.7 (ESRI, USA) to produce maps and run analyses. Maps use the Albers Equal Area Conic projection.

## 3 Results

Fourteen out of 20 contiguous US ecoregions have >1% of their area in deciduous or mixed forest, where deciduous forests are the majority ecosystem for two ecoregions (Table 1). The ecoregions with the greatest proportion of deciduous forest cover include the Atlantic Highlands in New England (62.0%), the Ozark/Appalachian forests (61.0%), and the mixed wood shield and plains in the upper Midwest (36.9%) (Figure 1a). Almost all ecoregions saw a steady decline in deciduous forest cover between 1985 and 2016, with exception of two prairie ecoregions and the central plains. In general, ecoregions with less forest cover saw greater percent change in deciduous forest area in the past two decades (logarithmic relationship, r^2^ = 0.14) with highest percentage changes in the upper Gila Mountains (−82.2%), Tamaulipas (Texas) semi-arid plains (−79.9%), and Western Sierra Madre Piedmont (−70.8%) (Figure 1a). Overall, changes ranged from −82.2% to +2.25% for ecoregions and totaled −17.5% for the contiguous US.

**Table 1.** Percent deciduous/mixed forest cover from 1985-2050 for ecoregions in the contiguous United States. Percent of the ecoregion that falls within protected areas managed consistently with biodiversity conservation (USGS PADUS, 2018; GAP code 1 or 2) is included. For projected forest coverage, 2050a represents the economic-growth scenario and 2050b represents the sustainability scenario.

| Ecoregion | Area | % Deciduous/Mixed Forest Cover |  |  |  |  |  |  |  |  |  |  |  | %<br>Protecte<br>d |
| --- | --- | --- | --- | --- | --- | --- | --- | --- | --- | --- | --- | --- | --- | --- |
| Level II | (ha) | 1985 | 1988 | 1991 | 2001 | 2004 | 2006 | 2008 | 2011 | 2013 | 2016 | 2050<br>a | 2050<br>b | (PADUS) |
| Atlantic Highlands | 1.51E+07 | 68.64 | 68.89 | 69.07 | 62.01 | 63.28 | 62.82 | 62.59 | 62.23 | 62.27 | 62.02 | 63.42 | 68.24 | 13.00 |
| Ozark/Appalachian Forest | 5.21E+07 | 65.12 | 65.18 | 65.16 | 60.12 | 61.39 | 60.93 | 60.87 | 60.68 | 60.93 | 60.97 | 57.57 | 63.75 | 8.61 |
| Mixed Wood Shield | 2.14E+07 | 43.29 | 43.52 | 43.76 | 38.72 | 37.08 | 36.86 | 36.95 | 37.00 | 37.11 | 36.94 | 38.14 | 42.22 | 22.85 |
| SE Plains | 1.03E+08 | 36.96 | 37.28 | 37.50 | 27.19 | 26.60 | 26.26 | 26.02 | 25.81 | 26.00 | 25.86 | 30.54 | 38.69 | 2.47 |
| Mixed Wood Plains | 3.92E+07 | 36.37 | 36.72 | 37.09 | 31.68 | 32.82 | 32.67 | 32.63 | 32.52 | 32.54 | 32.45 | 31.15 | 38.36 | 2.94 |
| Central Plains | 2.26E+07 | 8.37 | 8.68 | 9.24 | 9.90 | 10.04 | 9.98 | 9.98 | 9.95 | 9.95 | 9.93 | 7.59 | 9.59 | 1.53 |
| SE Coastal Plains | 3.40E+07 | 7.65 | 7.69 | 7.64 | 2.78 | 2.64 | 2.59 | 2.60 | 2.64 | 2.66 | 2.66 | 6.42 | 7.80 | 8.73 |
| Mediterranean CA | 1.64E+07 | 7.36 | 7.38 | 7.38 | 5.29 | 4.50 | 4.44 | 4.14 | 4.13 | 4.13 | 3.99 | 7.40 | 7.43 | 10.25 |
| TX/LA Coastal Plain | 7.36E+06 | 7.32 | 7.36 | 7.35 | 3.00 | 3.03 | 2.97 | 2.95 | 3.00 | 3.04 | 3.03 | 6.33 | 9.49 | 5.23 |
| Marine West Coast | 3.64E+07 | 6.18 | 6.19 | 6.19 | 3.35 | 3.06 | 2.99 | 2.97 | 2.99 | 3.02 | 3.05 | 5.86 | 6.14 | 63.33 |
| Temperate Prairies | 5.22E+07 | 6.16 | 6.17 | 6.18 | 6.01 | 6.58 | 6.55 | 6.54 | 6.53 | 6.54 | 6.55 | 5.61 | 6.15 | 2.37 |
| Western Cordillera | 8.22E+07 | 6.04 | 6.04 | 6.03 | 4.95 | 4.61 | 4.58 | 4.57 | 4.57 | 4.58 | 4.60 | 5.93 | 5.93 | 22.70 |
| Semiarid Plain | 5.31E+06 | 3.22 | 3.20 | 3.20 | 0.61 | 0.65 | 0.65 | 0.64 | 0.65 | 0.65 | 0.65 | 2.75 | 3.03 | 0.32 |
| South Central Prairies | 9.97E+07 | 3.21 | 3.19 | 3.16 | 3.32 | 3.35 | 3.31 | 3.29 | 3.26 | 3.26 | 3.28 | 2.73 | 3.08 | 0.88 |
| Upper Gila Mtns | 1.09E+07 | 1.00 | 1.00 | 0.99 | 0.21 | 0.22 | 0.22 | 0.22 | 0.21 | 0.18 | 0.18 | 0.97 | 0.97 | 11.24 |
| West-Central Prairies | 5.91E+07 | 0.79 | 0.79 | 0.79 | 0.59 | 0.57 | 0.57 | 0.57 | 0.57 | 0.57 | 0.57 | 0.79 | 0.80 | 2.43 |
| Western Sierra<br>Madres | 4.31E+0<br>6 | 0.70 | 0.70 | 0.70 | 0.28 | 0.28 | 0.28 | 0.28 | 0.20 | 0.20 | 0.20 | 0.70 | 0.70 | 12.97 |
| Cold Deserts | 1.01E+0<br>8 | 0.65 | 0.65 | 0.65 | 0.58 | 0.56 | 0.55 | 0.55 | 0.55 | 0.54 | 0.55 | 0.65 | 0.65 | 12.45 |
| Warm Deserts | 4.08E+0<br>7 | 0.14 | 0.14 | 0.14 | 0.02 | 0.12 | 0.12 | 0.12 | 0.09 | 0.09 | 0.09 | 0.14 | 0.14 | 36.62 |
| Everglades | 2.20E+0<br>6 | 0.00 | 0.00 | 0.00 | 0.07 | 0.37 | 0.36 | 0.36 | 0.36 | 0.36 | 0.36 | 0.00 | 0.00 | 38.06 |

**Fig. 1.**
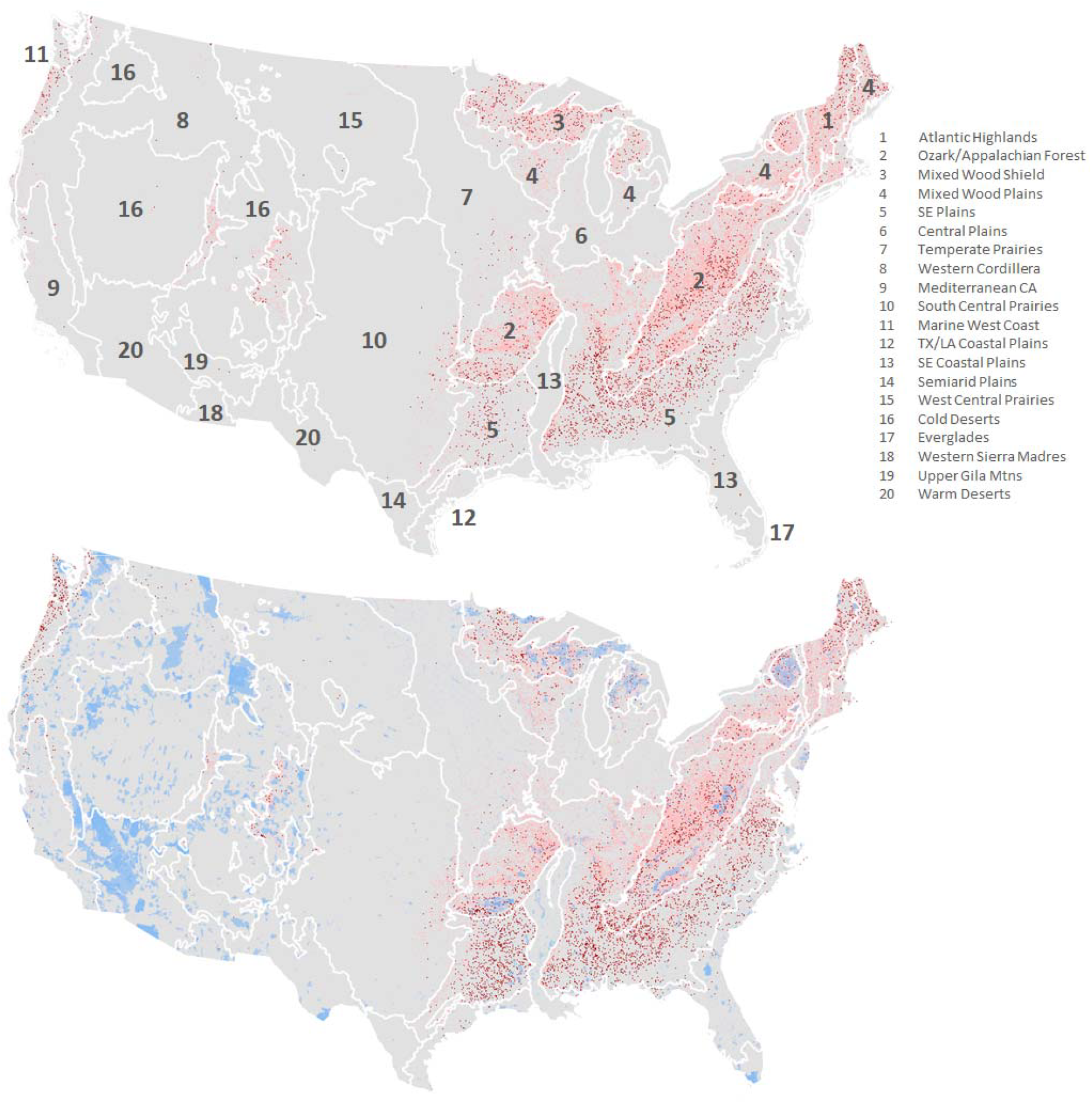
Ecoregions of the contiguous United State with deciduous/mixed forest cover (pink) and disturbed forest (red) from a) 1985-2016 and b) models projecting forest cover in 2050. Numbers indicate rankings of ecoregions by proportion of forest cover in 2016. Blue areas represent protected areas managed consistently with biodiversity conservation (USGS PADUS GAP code 1 or 2).

Predictions of deciduous forest cover depended heavily on the scenario employed. The “economic growth scenario” resulted in higher rates of forest loss due to urban increase, agricultural expansion, and higher demand for forest products. Most ecoregions remain in decline by 2050 with the exception of deserts, Mediterranean California and Western Sierra Madre Piedmont. For the top leaf-peeping ecoregions, predicted percent change in forest cover is nearly equivalent to declines from 1985-2016 (Figure 1b). Under the “sustainability scenario,” more than half of the ecoregions would have increasing forest cover by 2050. Generally, highest percent increases would occur in ecoregions with relatively low forest cover (logarithmic relationship, r^2^ = 0.12).

Ecoregions for which deciduous forests make up at least a quarter of the land cover have anywhere between 2.5% and 22.8% protected areas coverage. Declines in forest cover between 1985 and 2050 (economic-growth scenario) exhibited a logarithmic relationship with protected areas coverage across ecoregions, with ecoregions undergoing greater proportional losses having greater forest area and being more often underrepresented in the protected areas network (Figure 2; r^2^ = 0.35, *p* = 0.03). Under the “sustainability scenario,” ecoregions with higher forest cover, but low representation in the protected areas network (i.e., mixed wood plains and southeast plains) will see relatively higher gains in deciduous forest.

**Fig. 2.**
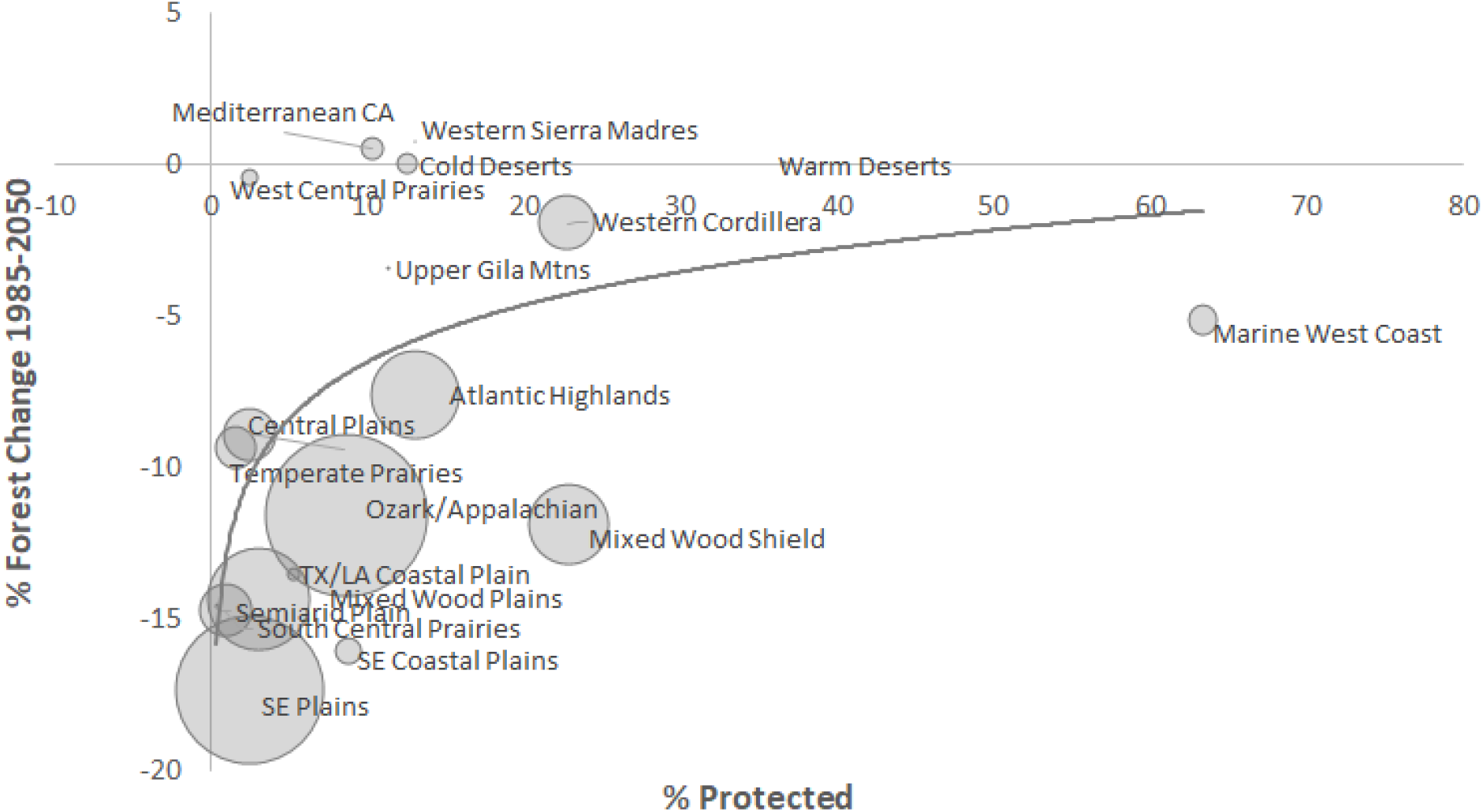
Percent change in deciduous/mixed forest cover in an ecoregion from 1985 to projections for 2050 as a function of forested area and the percent of the ecoregion that is protected as of 2018 (USGS PADUS GAP code 1 or 2; y = 0.14(protected) - 2.9E-11(forest) - 8.01, r^2^ = 0.35, *p* = 0.03).

## 5 Conclusions

Forests are an essential component of the natural environment that offer profound aesthetic and spiritual benefits to humanity. As such, forests offer a unique and powerful way to connect people with environmental issues like the biodiversity and climate crises (Ives et al., 2018). Loss of forests, either in large tracts or piecemeal, results in a degradation of these benefits and decreases opportunities for quality experiences in nature (Nisbet et al., 2008; Soga and Gaston, 2016; Zylstra et al., 2014). Here we documented extensive temperate forest loss through the context of fall color change. These losses may be changing how people experience nature, such as through the awe inspired by leaf peeping.

Cumulatively, we found that just under 55 million acres of deciduous or mixed forest were disturbed between 1985 and 2016. All ecoregions with deciduous forests incurred losses, but some were more dramatic than others. As expected, the extent of loss was related to the proportion of the ecoregion that is managed as multi-use or is lacking conservation mandates. Underrepresentation in the U.S. protected areas network means that temperate forests are susceptible to continued fragmentation and modification. Under worst-case forecasting scenarios, losses are predicted to continue. However, environmentally focused scenarios suggest there is still opportunity to reverse deciduous forest loss in some ecoregions. Increasing public exposure to temperate forests may help ensure conservation of more natural areas and preserve the quantity and quality of autumn forest viewing.

The top ecoregions for autumn aesthetics are experiencing relatively higher forest losses and are also relatively under-protected. Collectively, ecoregions with greater than a quarter of their area covered by deciduous/mixed forest saw a 16.5% decline in forest area. For these ecoregions, 93% of their lands are currently unprotected suggesting they may face further disturbance or fragmentation. The top ecoregions are also concentrated east of the Mississippi River, where there is less public land and more human disturbance, elevating the importance of restoration efforts (Jenkins et al., 2015; Rosa and Malcom, 2020). Spatial incongruities in protected areas and forested ecoregions also impact other ecosystem values. For example, previous research indicates that the siting for current protected areas is often discordant with diversity-rich or carbon-rich areas of the country (Jenkins et al., 2015; Rosa and Malcom, 2020). This means the remaining natural habitats in these areas are significantly under-protected; elevating their protections could contribute to both climate mitigation and biodiversity conservation. In particular, states across the Southeast harbor high levels of biodiversity and very few protected areas.

Additionally, trends show that ecoregions with lower forest cover generally experienced greater rates of decline in the past two decades; this suggests that the few cultural and habitat resources that deciduous forests provide in these areas are dwindling relatively fast. In the United States, conversion of privately owned rural lands into low-density residential development (i.e., exurban development) has increased five- to seven-fold between 1950 and 2000 and is the fastest growing type of land use (Suarez-Rubio et al., 2013). Exurban development is prominent in forested ecoregions with dispersed, isolated housing units embedded within a forest matrix. At a more local scale, topography will likely remain a significant constraining factor in development, allowing some areas to persist in forest cover while agricultural, residential and urban uses are concentrated in specific portions of the landscape (Wear and Bolstad, 1998). However, land use may intensify without associated changes in land cover if development occurs under the forest canopy: in some cases, forest cover is increasing (rather than declining) with an increase in human population density and development. This suggests that estimates of forest cover and ecosystem impacts derived from satellite imagery are conservative and presents limitations to continued tracking and prediction of future trends in forest disturbance.

Projected land use patterns suggest spatial variability in continued anthropogenic landscape modifications. Some ecoregions, mostly prairies and plains, may continue to see forest losses regardless of scenario. But for some, like the mixed wood plains (32.5% deciduous/mixed forest cover), which covers regions of the northeastern coast and upper Midwest, human behaviors could mean the difference between continued forest losses or reversal to gains. Models under the sustainability scenario suggest there is still time to reverse declining trends in deciduous forest loss; 21.5% increase in forest cover is projected for the top five forested ecoregions, collectively. However, even if forest loss is projected to slow or reverse, this may not directly equate to impacts to cultural ecosystem values because of lag times in forest responses to stress (Alexander et al., 2018). Additionally, these are net changes and do not capture spatial changes at a local level where the loss and gain of forest may be disjunct, ultimately resulting in more cumulative forest area, but also more fragmented forest patches. Ecoregions that lack protections will continue to experience greater forest disturbances, which is consistent with global analyses (d’Annunzio et al., 2015). These disturbances (e.g., exurban developments) are often close to protected areas and natural amenities raising concern about their ecological consequences, this is one of many stressors that deciduous forests will continue to face.

Centuries of exploitation demonstrate the resilience of temperate forests, but how much disturbance can be tolerated and what it may mean for forest health and fall colors is still under investigation. Currently, the majority of science on autumn colors focuses on the variation of color pigments and senescence timing with nutrient availability, gene expression, herbivory, and climatic factors. Additionally, there are studies on the impacts of fragmentation and degradation on other forest values such as habitat, but any connections to be made between these stressors and autumn aesthetic will be inferential at best given the current lack of science focused on direct relationships. Understanding the indirect impacts of exurban development and forest fragmentation on fall colors could be one way to generally quantify impacts to cultural ecosystem values without the extra complication of estimating monetary values. For example, under future climate change projections, greater summer heat-stress will cause abbreviated leaf coloration seasons for most tree species (Xie et al., 2018). However, temperature discrepancies which would typically be exacerbated by fragmentation may be dampened by the obscured edges between exurban and forest land covers, highlighting the importance of locall□ and landscapel□scale features on microclimate heterogeneity (Arroyo-Rodriguez et al., 2017; Latimer and Zuckerberg, 2016). In addition to landscape conversion, forests are experiencing increasing frequency, extent, and severity of natural disturbances, anthropogenic climate change, and a burgeoning global human population that imposes escalating demands on forests. However, the current body of literature fails to present a strong understanding of their synergistic impacts. For example, stressed trees may senescence earlier than others, but research also suggests that climate warming can delay timing (Faticov et al., 2019; Schaberg et al., 2003). Research demonstrates increased vulnerability of temperate forests to disease and invasion during periods of stress, but impacts to autumn aesthetics are unknown. More research is needed to determine how forest loss and fragmentation relate to spatiotemporal changes in autumn color vibrancy and to the quality of human-nature connections.

The large difference in forest loss estimates in the predictions scenarios emphasizes the importance of human approaches to economic growth and sustainability in securing environmental stability. Scenic aesthetic is the most direct and immediate aspect via which people perceive and begin to value landscapes (Gobster et al., 2007; Sargolini, 2013). Visually appealing and healthy ecological landscapes evoke positive emotions and promote the desire to protect such landscapes (Gobster et al., 2007; Lee, 2017). Growing opportunities for U.S. conservation could work to forge deeper connections between humans and nature and motivate the public to take protective actions against detrimental environmental changes. In response to current global biodiversity and climate crises, science-driven guidance to protect 30% of global lands and seas by 2030 have made its way into US federal and state policy proposals. These proposals call for achieving more equitable access to public land, nature and a healthy environment for all communities. Given this framework, the potential benefits of protecting deciduous forests is manifold. In addition to preserving the visual aesthetics and spiritual connections that people make during autumn senescence, conserving temperate forests also means protecting many of the U.S.’s biodiversity hotspots and areas of high carbon potential. Additionally, forest conservation ensures more ecosystems will be more resilient to climate stresses (Xu et al., 2019). Therefore, encouraging the public to experience temperate forest autumn foliage may in turn have broad reaching conservation implications for the future.

## Acknowledgments, Samples, and Data

We thank T. Niederman for her thoughtful review of this manuscript. We also thank USGS for making PADUS, MRLC for making NLCD, and T. Sohl et al. for making their datasets available to streamline general analyses. The authors received no additional financial support for the research, authorship and/or publication of this article. The authors declare that the research was conducted in the absence of any commercial or financial relationships that could be construed as a potential conflict of interest. Analyses reported in this article can be reproduced using publicly available data. Final outputs are available on OSF at https://osf.io/bqn7k/.

